# In vivo Targeting MEK and TNK2/SRC pathways in PTPN11 driven leukemia

**DOI:** 10.1101/2024.05.16.594555

**Authors:** Bill H. Chang, Karina Thiel-Klare, Jeffrey W. Tyner

**Author notes:** Correspondence to: Bill H Chang, MD, PhD, 3181 SW Sam Jackson Park Road, Portland, OR 97239.

## Abstract

PTPN11 encodes for a tyrosine phosphatase implicated in the pathogenesis of hematologic malignancies such as Juvenile Myelomonocytic Leukemia (JMML), Acute Myeloid Leukemia (AML), and Acute Lymphoblastic Leukemia (ALL). Since activating mutations of PTPN11 increase proliferative signaling and cell survival through the RAS/MAPK proliferative pathway there is significant interest in using MEK inhibitors for clinical benefit. Yet, single agent clinical activity has been minimal. Previously, we showed that PTPN11 is further activated by upstream tyrosine kinases TNK2/SRC, and that PTPN11-mutant JMML and AML cells are sensitive to TNK2 inhibition using dasatinib. In order to validate these findings, we adopted a genetically engineered mouse model of PTPN11 driven leukemia using the mouse strain 129S/Sv-*Ptpn11*^*tm6Bgn*^/Mmucd crossed with B6.129P2-*Lyz2*^*tm1(cre)Ifo*^/J. The F1 progeny expressing Ptpn11^D61Y^ within hematopoietic cells destined along the granulocyte-monocyte progenitor lineage developed a fatal myeloproliferative disorder characterized by neutrophilia and monocytosis, and infiltration of myeloid cells into the liver and spleen. Cohorts of Ptpn11^D61Y^ expressing animals treated with combination of dasatinib and trametinib for an extended period of time was well tolerated and had a significant effect in mitigating disease parameters compared to single agents. Finally, a primary patient-derived xenograft model using a myeloid leukemia with PTPN11 F71L also displayed improved disease response to combination. Collectively, these studies point to combined therapies targeting MEK and TNK2/SRC as a promising therapeutic potential for PTPN11-mutant leukemias.

**Key Points:**

- Combining MEK and TNK2/SRC inhibitors has therapeutic potential in PTPN11 mutant JMML and AML

## Introduction

PTPN11 is a ubiquitous protein tyrosine phosphatase that activates many signaling pathways including RAS/MAPK(1). Disease-associated gain-of-function mutations in PTPN11 have been identified as the most common driver of juvenile myelomonocytic leukemia (JMML), found in 35% of cases(2, 3). Similar mutations in PTPN11 are present in ∼5-10% of acute myeloid leukemia (AML) cases, and ∼5% of B-cell acute lymphoblastic leukemia (ALL)(2-7). While in other cancers, mutations in RAS/MAPK pathway may be a secondary event, JMML is driven by initiating mutations in the RAS/MAPK pathway(3, 8), making this disease an ideal model for targeting this pathway for treatment. JMML remains a difficult to treat disease with allogeneic stem cell transplant as the only known cure with a 50% failure rate at 2 years post-transplant(9). These current clinical outcomes make development of new therapies imperative.

Recent studies targeting RAS/MAPK activity through MEK inhibition have been validated in multiple diseases(10). Further, preclinical studies have validated MEK inhibition in models of JMML and in patient derived xenografts (11, 12). This has led to a national phase II study of relapsed JMML using trametinib, a potent and selective MEK inhibitor (NCT03190915, ADVL1521)(13). Although responses were seen, alternate approaches appear to be necessary. Epigenetic signatures have further been hypothesized to represent another mechanism underpinning the pathogenesis of JMML(14-17) leading to a strong rationale for combining trametinib with hypomethylating agents(12) in the ongoing clinical trial (NCT05849662, TACL2020-004).

We previously described a patient with refractory JMML who was treated as a single agent with the multikinase inhibitor, dasatinib(18). Molecular studies identified a novel mechanism of action whereby Tyrosine Kinase Non-receptor 2 (TNK2) increases activity and dependence in PTPN11 mutated leukemias(18). TNK2 is a cytoplasmic tyrosine kinase that is overexpressed in many solid tumors(19). Although no current TNK2 selective inhibitor has been developed for clinical use, the multikinase inhibitor dasatinib has been shown to inhibit TNK2(20). As proof of concept our patient with PTPN11 mutant JMML had a clinical response to dasatinib suggesting targeting TNK2 activity may have therapeutic effect in PTPN11 mutant driven leukemias. Further, other studies suggested a potential therapeutic effect for dasatinib in regulating SRC activity in the proliferation signal in JMML(21).

Although single agent may have therapeutic effect, it does appear to be transient providing the basis for further combination therapy. Thus, we proceeded to test whether combination of dasatinib with trametinib may improve therapeutic effect in PTPN11 mutant driven leukemias. To test this mechanism, we adopted a mouse model of PTPN11 driven JMML. A transgenic mouse model by Neel *et al* was developed as a conditional knock-in of Ptpn11^D61Y^ in the mouse strain 129S/Sv-*Ptpn11*^*tm6Bgn*^/Mmucd (carrying a *loxP*-neo-*loxP*-STOP-*loxP* cassette between exons 2 and 3 of *Ptpn11* and inserting a D61Y point mutation)(22). In order to control postnatal expression of this allele, the authors initially used Mx1-Cre to induce knock-in at 3-4weeks of age using polyinosinic-polycytidylic acid. These animals developed a fatal myeloproliferative disorder within 6 months consistent with JMML. With some concern for the Mx1-Cre system(23) we chose to cross this strain B6.129P2-*Lyz2*^*tm1(cre)Ifo*^/J(24) expressing Cre in myeloid cells at the M lysozyme locus to produce a progeny that express Ptpn11^D61Y^ mostly within myeloid hematopoietic progenitor cells. Herein, we describe the *in vivo* effects using this Ptpn11^D61Y^ mouse model as well as a primary AML patient-derived xenograft treated with prolonged exposure to combination of dasatinib and trametinib with potential benefit.

## Material and Methods

### Patient Samples

All clinical specimens were collected with informed consent on a protocol approved by the Oregon Health & Science University (OHSU) Institutional Review Board (IRB #4422).

### Transgenic mouse colonies

All murine studies were performed in accordance with OHSU IACUC (IP00000179), maintained in barrier conditions and monitored through the Department of Comparative Medicine at OHSU. We obtained the mouse strain 129S/Sv-*Ptpn11*^*tm6Bgn*^/Mmucd (MMRRC at UC Davis) that carries a *loxP*-neo-*loxP*-STOP-*loxP* cassette between exons 2 and 3 of *Ptpn11* and inserts a D61Y point mutation(22). These animals were crossed with B6.129P2-*Lyz2*^*tm1(cre)Ifo*^/J(24) (Jackson Laboratories) that express the Cre recombinase under the control of the endogenous *Lyz2* promoter/enhancer elements. Thereby, the F1 progeny express Ptpn11^D61Y^ within hematopoietic cells destined along the granulocyte-monocyte progenitor lineage. Validation of the F1 progeny genotype was performed as previously described through TransnetYX(22). Animals were monitored monthly with peripheral blood counts using the Element HT5 veterinary hematology analyzer (Heska). NOD.*Cg-Prkdc*^*scid*^ *Il2rg*^*tm1Wjl*^Tg(CMVIL3,CSF2,KITLG) 1Eav/MloySzJ (NSGS)(25) (Jackson Laboratories) mice were used for primary AML xenograft experiments.

### AML patient derived Xenograft

Six-week-old NSGS females were sub-lethally irradiated with 150cGy followed 24 hours later by tail vein injection of approximately 0.6×10^6 mononuclear cells per animal. The AML sample underwent human CD3 depletion using dynabead CD3 depletion kit (Invitrogen) prior to injection. Animals were monitored for engraftment starting 4 weeks from injection with blood draws. Peripheral blood was monitored for engraftment by flow cytometry using a cocktail of human CD45-FITC (Milteny), human CD33-APC (BD Pharmingen), human CD3-PE (Invitrogen), and mouse CD45-erpCP Cy5.5 (BD Biosciences) and analyzed on an LSRFortessa at the OHSU Knight Cancer Institute Flow Cytometry Core. Engraftment was determined when animals had approximately 1% human CD33/human CD45 chimerism prior to initiation of treatment.

### *In vivo* drug treatment

Drug dosing was performed as previously described for dasatinib (40mg/kg/dose(26)) and trametinib (1mg/kg/dose(27)). Dasatinib (LC labs) was compounded in 80nM Sodium Citrate, pH 3.1 at 8mg/ml while trametinib (LC labs) was first dissolved in dimethylsulfoxide (DMSO) at 40mg/ml and then compounded in 0.5% hydroxypropylmethylcellulose/0.2% Tween 80 to a final concentration of 0.2mg/ml. Animals were treated once daily by oral gavage either by vehicle or by drug and were monitored for disease by peripheral blood counts as well as behavior and well-being.

### *Ex vivo* inhibitor Assay

*Ex vivo* assessment of drug response as single agent and combination was performed as previously described(28). Briefly, small-molecule inhibitors were purchased from LC Laboratories (Woburn, MA), or Selleck Chemicals (Houston, TX), and reconstituted in DMSO and stored at −80 °C. Inhibitors were distributed into 384-well plates prepared with each single-agent in a seven-point, 3-fold concentration series ranging from 0.0137 μM to 10 μM as well as combined in an equimolar series of the same 7 concentrations. The final concentration of DMSO was ≤0.1% in all wells. Freshly isolated mononuclear cells were seeded into prepared drug plates at 1×10^4 cells per well in RPMI 1640 media supplemented with FBS (20%), L-glutamine, and penicillin/streptomycin. At the time of seeding, cell viability was >90%. After 3 days of culture at 37 °C in 5% CO_2_, MTS reagent (CellTiter96; Promega, Madison, WI) was added, optical density was measured at 490 nm, and raw absorbance values were blanked and normalized to untreated control wells to determine relative cell viability. A mixture of flavopiridol, staurosporine, and velcade (F-S-V) was used as a positive control treatment well. Normalized viability data were fit to a probit regression curve to determine IC_50_.

### Colony formation assays

Bone marrow was harvested from mice by flushing the femur and tibia with media. Approximately 1×10^5 cell per ml were plated in triplicate into M3234 incomplete methylcellulose medium (StemCell Technologies) supplemented with 0.1 ng/mL GM-CSF (Peprotech). Cells were visualized at day 7 or 14 using the STEMVision colony counter (STEMCELL Technologies) with manual counting to ensure accuracy.

### Statistical analysis

Mean values ± SEM are shown unless otherwise stated. One-way ANOVA or two-tailed student’s *t*-test were used for comparisons. Statistical analysis was conducted using GraphPad Prism Version 6.0. Sample sizes of approximately five animals per cohort were consistent with those reported in similar studies and provide sufficient power to detect changes with the appropriate statistical analysis.

## Results

### Ptpn11^D61Y^ animals develop a myeloproliferative neoplasm

129S/Sv-*Ptpn11*^*tm6Bgn*^/Mmucd crossed with B6.129P2-*Lyz2*^*tm1(cre)Ifo*^/J produced healthy pups with approximately twenty-five percent of the litter expressing Ptpn11^D61Y^. Consistent with the original characterization of the Ptpn11^D61Y^ strain(22), the Ptpn11^D61Y^ progeny eventually develop a myeloproliferative disorder within eight months of age (Figure 1A). This disorder was manifested by quite pronounced splenomegaly greater than 10x larger than WT and hepatomegaly 3x larger than WT (average control spleen weight of 85mg compared to Ptpn11 spleen weight of 2298mg, p=0.0002, Figure 1B). Peripheral blood examination identified a leukocytosis with absolute neutrophilia and monocytosis. Absolute lymphocytosis was also seen although was decreased relative to the other cell lineages (Figure 1C). Although hemoglobin and platelets were both decreased the mean platelet volume and polychromasia on the smear would suggest a consumptive process (Figure 1D). Taken together, these findings support this animal model is consistent with Ptpn11-driven myeloproliferation.

**Figure 1.**
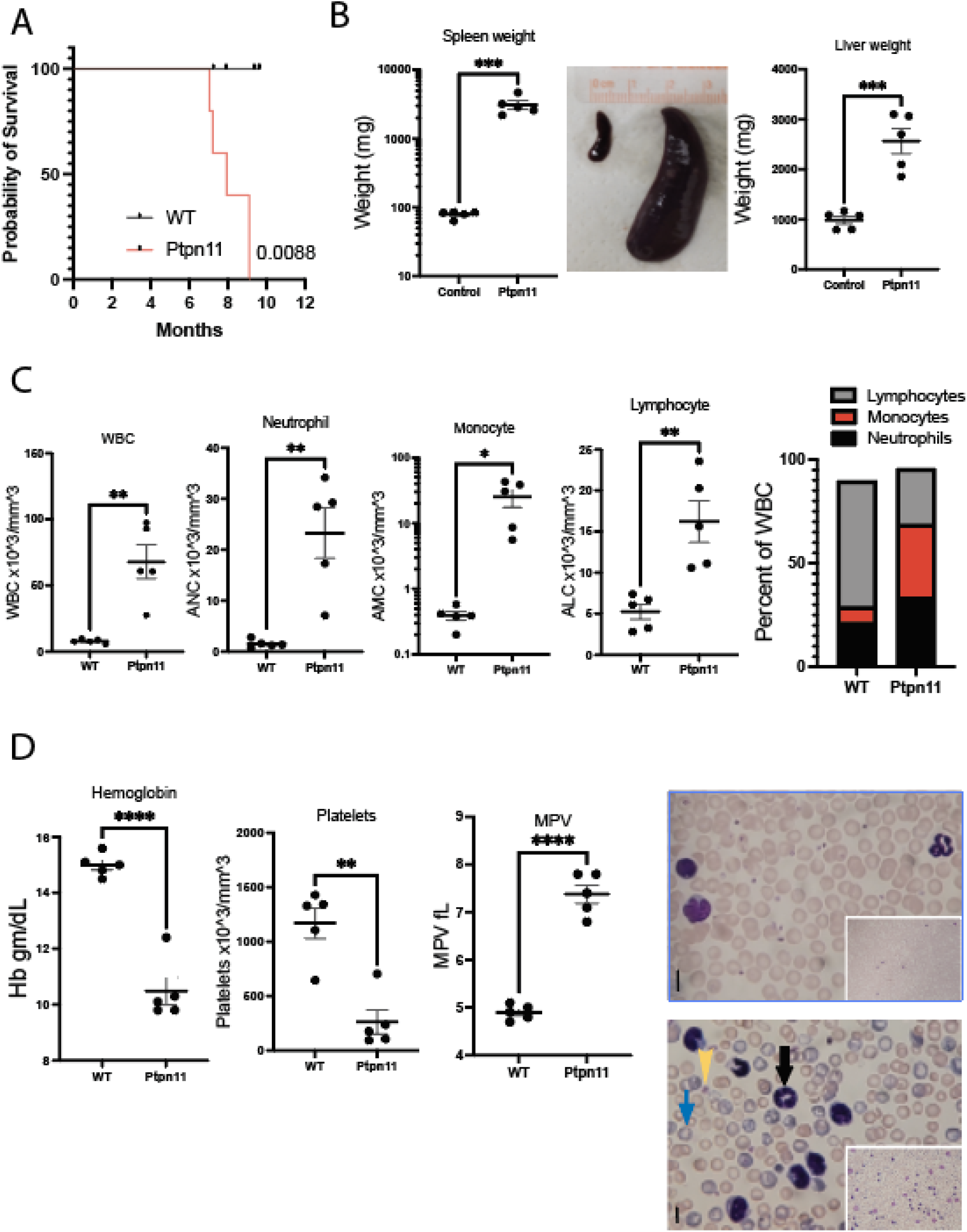
Ptpn11^D61Y^ mice develop a myeloproliferative disorder. **A)** Survival of cohorts of Ptpn11 wildtype (WT) and Ptpn11^D61Y^ (Ptpn11) littermates. A cohort of five Ptpn11 animals and five age matched WT counterparts were followed over time. Ptpn11 animals developed signs of disease within 8 months compared to their WT counterparts (p=0.0088, Log-Rank). **B)** Spleen and liver weights at sacrifice of Ptpn11 animals compared to their age matched WT counterparts. The average spleen weight of Ptpn11 animals was 2300mg compared to WT animals of 85mg (p=0.0002***) with representative image. The average liver weight of Ptpn11 animals was 2500mg compared to WT animals of 980mg (p=0.0003***). **C)** Peripheral blood counts, including white blood cell counts (WBC), neutrophils, monocytes, and lymphocytes. The total WBC (mean 67.8 v 7.9 ×10^3/mm^3, p=0.0015**), absolute neutrophil (23.3 v 1.5 ×10^3/mm^3, p=0.0022**), monocyte (25.3 v 0.4 ×10^3/mm^3, p=0.012*), and lymphocyte (16.2 v 5.5 ×10^3/mm^3, p=0.0036**) counts were elevated in the Ptpn11 animals. **D)** Hemoglobin and platelets at the time of harvest. Ptpn11 animal showed a decrease in hemoglobin (10 v 15 gm/dL, p<0.0001****) and platelets (263 v 1200 ×10^3/mm^3, p=0.0011**) with an elevation in mean platelet volume (MPV) (7.4 v 4.9 fL, p<0.0001). Photomicrographs of GIEMSA stained peripheral blood at 10x (inset) and 40x power. Upper panel of WT blood compared to lower panel of Ptpn11. The peripheral smear of Ptpn11 animals display increased WBC including neutrophils (black arrow), RBC’s consistent with polychromasia (blue arrow), and a decrease in platelets (yellow arrow), suggestive of a consumptive process. Black line denotes 0.01mm. Images were captured with a Leica DMLS microscope and captured on Axiocam digital camera. Statistical comparisons were performed with a student’s t-test.

### Ex vivo *response to MEK and TNK2/SRC inhibition*

Increased hypersensitivity of hematopoietic cells to the cytokine GM-CSF in myeloid colony formation assays is a hallmark of JMML pathogenesis(1, 29). To test the effects of this hypersensitivity, mononuclear cells extracted from the bone marrow of Ptpn11^D61Y^ animals were harvested at 9months of age and plated onto cytokine free methylcellulose with GM-CSF (Figure 2A). Separate wells were further exposed to dasatinib, trametinib, or equimolar combination. After seven days in culture, the colonies were quantified. Trametinib and combination of drug showed a significant decrease in the size and number of colonies. Viability of mononuclear cells extracted from Ptpn11^D61Y^ spleens also showed a dose-response to both drugs with an additive effect in combination (Figure 2B). A matrix of combination was also performed to calculate a BLISS, HAS, Loewe, and ZIP score. All scores landed between -10 and 10 predicted to be an additive effect from the combination.

**Figure 2.**
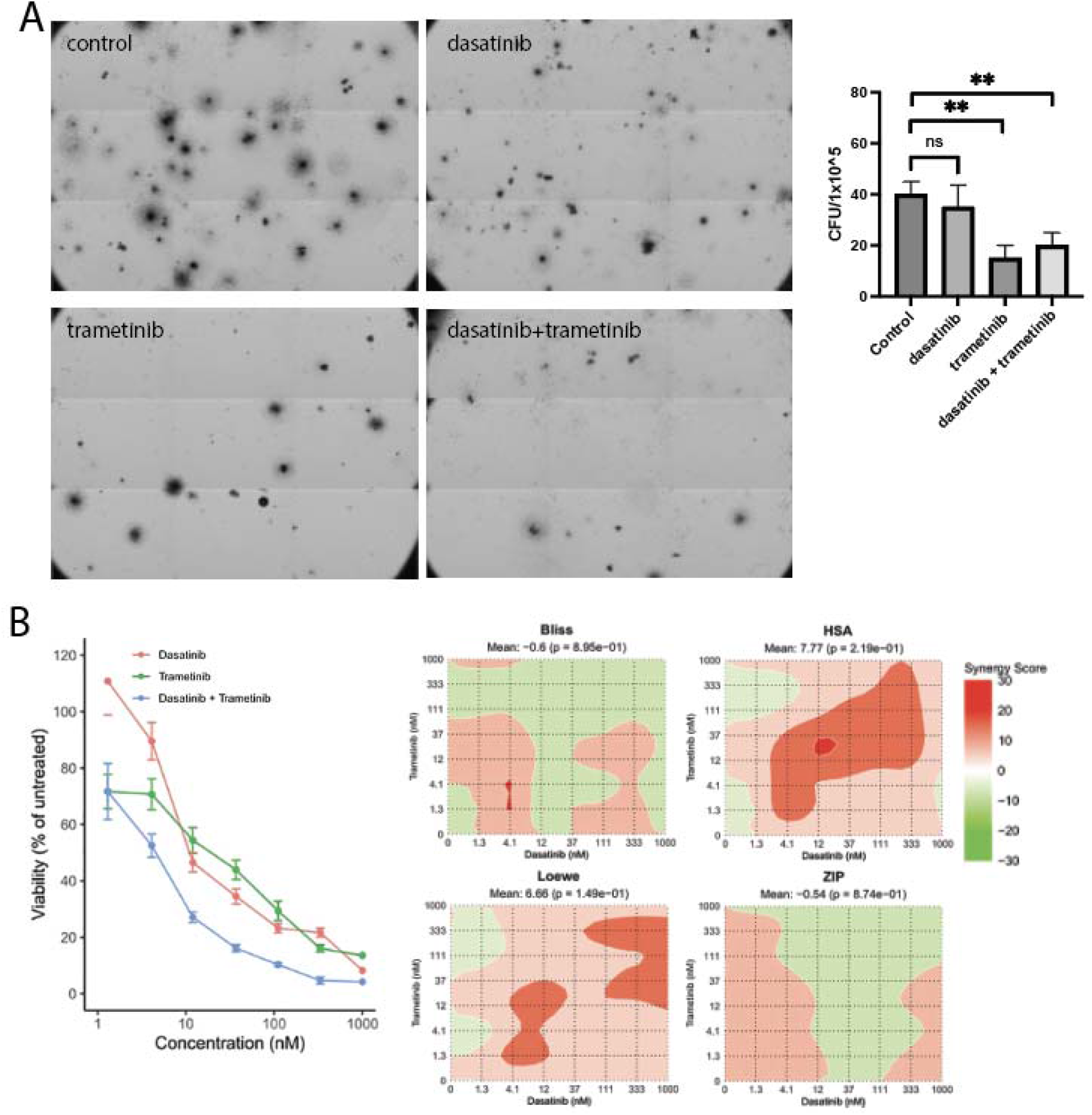
In vitro sensitivity of Ptpn11^D61Y^cells. **A)** Colony forming assays. Mononuclear cell were isolated from the bone marrow of Ptpn11^D61Y^ expressing animals. Cells were then incubated in cytokine free methocult for seven days with 0.1ng/ml GM-CSF, and also in th presence of 10nM dasatinib, 10nM trametinib, or the combination. B) Graphic representation of the number of colonies with the presence of GM-CSF. Statistical comparisons were performed with a student’s t-test. **p<0.01. C)Dose-Response curves or mononuclear cells harvested from Ptpn11^D61Y^ expressing cells. Mononuclear cells were isolated from the spleen and plated with increasing concentrations of trametinib (red), dasatinib (green) or equimolar combinations of trametinib and dasatinib (blue). Cells were exposed to drug for 3 days, then assessed for viability using the colorimetric MTS. Viability was then normalized to no drug control. A matrix of combination was also performed to calculate a BLISS, HAS, Loewe, and ZIP score.

### In vivo sensitivity to MEK and TNK2/SRC inhibition

To next test *in vivo* response to drugs, cohorts of Ptpn11^D61Y^ animals at 9 months of age began treatment with respective vehicles, dasatinib, trametinib, or combination daily five days per week by oral gavage for 3 months. The animals tolerated the treatment well with no change in weight, appearance, or activity. After three months of treatment, due to significant leukocytosis in the control group, the animals were sacrificed for disease evaluation. At the time of harvest there was a significant effect of the combination treatment in mitigating disease parameters compared to single agents (Figure 3). Although single agents significantly reduced the total WBC, neutrophils and monocytes the combination had the most dramatic effect on blood leukocytes (average WBC of 75.5×10^6 /mm^3 in control animals versus 6.1×10^6/mm^3 in combination treated animals, p=0.0009). Further, the hemoglobin (average control with 9.9gm/dL compared to combination with 14gm/dL, p<0.0001) and platelets (average control with 216×10^6/mm^3 compared to 1962×10^6/mm^3, p=0.006) showed a significant increase with values similar to their WT littermates. Finally, there was a significant effect on spleen size with the 3 months of treatment as compared to the control group with combination treated mice having spleen weights similar to WT (average control spleen weight of 1814mg compared to the combination treated of 72mg comparable to WT spleen size, p=0.0046).

**Figure 3.**
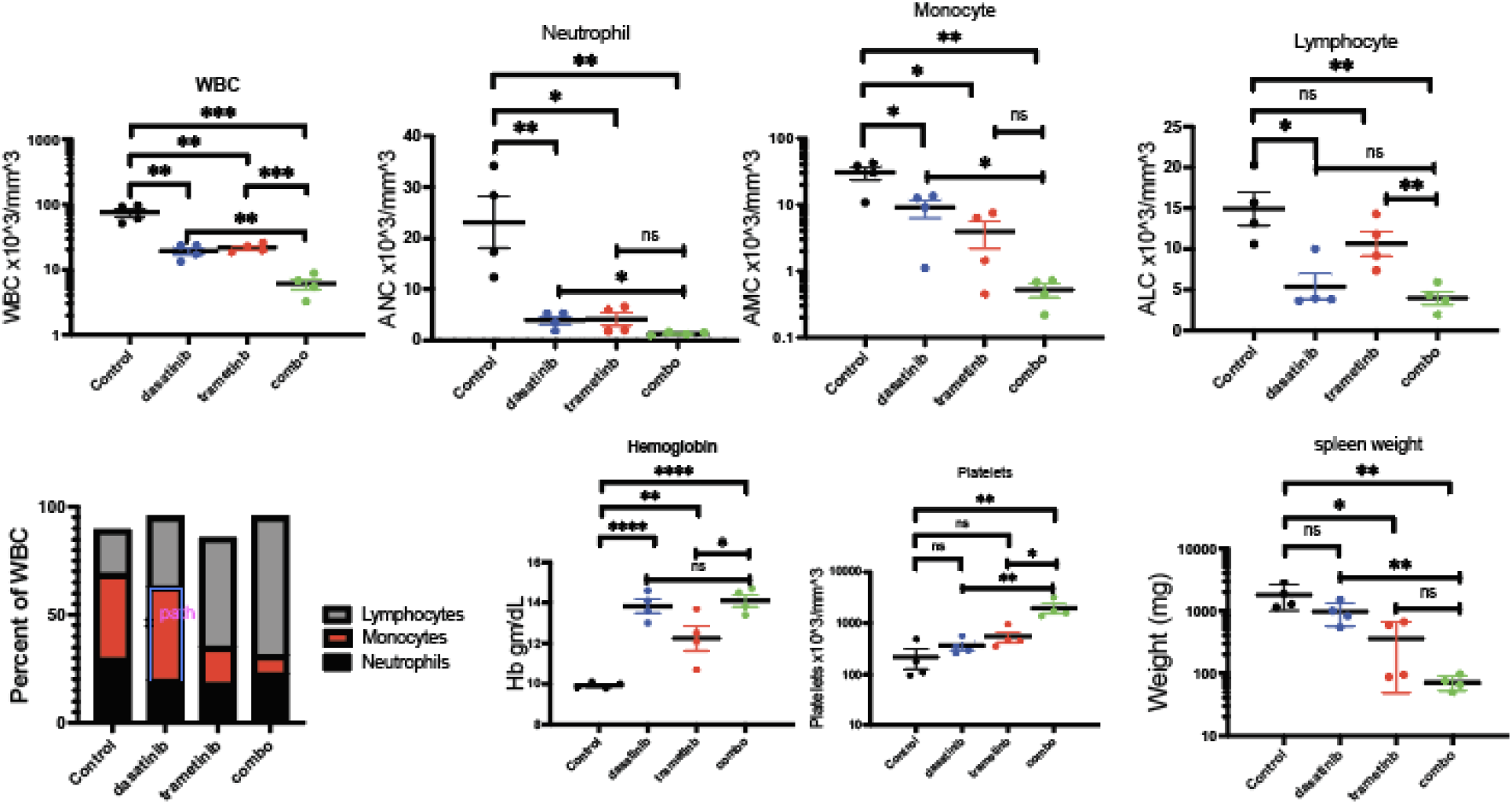
in vivo response to trametinib and dasatinib in Ptpn11^D61Y^ animals. Cohorts of five nine-month-old animals were treated with vehicle (Control), dasatinib (40mg/kg/day), trametinib (1mg/kg/day), or combination (40mg/kg/day dasatinib/1mg/kg/day trametinib). After 3 months of treatment, animals were sacrificed and assessed for disease status. Metrics included, blood counts with WBC, neutrophils, monocytes, lymphocytes, relative percentage of the total WBC, hemoglobin, platelets, and spleen weight at the time of harvest. Statistical comparisons were performed with a student’s t-test (*p<0.05, **p<0.01, ***p<0.001, p<0.0001).

Our previous studies suggest that PTPN11 mutant AML may also respond to combined inhibition(18). We currently have a biorepository (BEAT AML, local IRB4422) that has an extensive deposit of primary AML cells that have undergone genomic and functional interrogation(30, 31). Within this biorepository we identified a sample of acute monoblastic leukemia with normal cytogenetics. Molecular testing revealed npm p.W288fs*12, DNMT3A p.R882H, and PTPN11 p.P71L mutations. Functional interrogation of the sample through the BEAT AML biorepository program had already shown sensitivity to both trametinib and dasatinib as single agents (Figure 4A). This banked sample was thawed, T-cell lymphocyte depleted, injected into cohorts of NSGS mice and monitored for human chimerism in peripheral blood. Once animals showed some evidence of human chimerism of approximately 1 percent, cohorts began drug treatment daily for five days each week. Animals continued treatment until evidence of ≥25% human leukemic cells in the blood or moribund features. The study completed after 3 months of treatment and the remaining animals were harvested. At the end of 3 months or whenever animals met criteria for harvest, disease status was evaluated by spleen weight. There was a significant survival advantage in the animals treated in combination as compared to single agent or vehicle control (Figure 4B). It was further noted that at the end of 3 months, the animals treated with combination had a statistically improved disease status with the combination (Figure 4C) (average control spleen weight of 211mg compared to combination treated of 98mg, p=0.0417). Mononuclear cells extracted from the spleen continued to show some sensitivity to the drugs and an additive effect in combination (Figure 4D). Although the therapeutic effect appeared less dramatic in this AML model it continued to validate this combination as a potential therapy.

**Figure 4.**
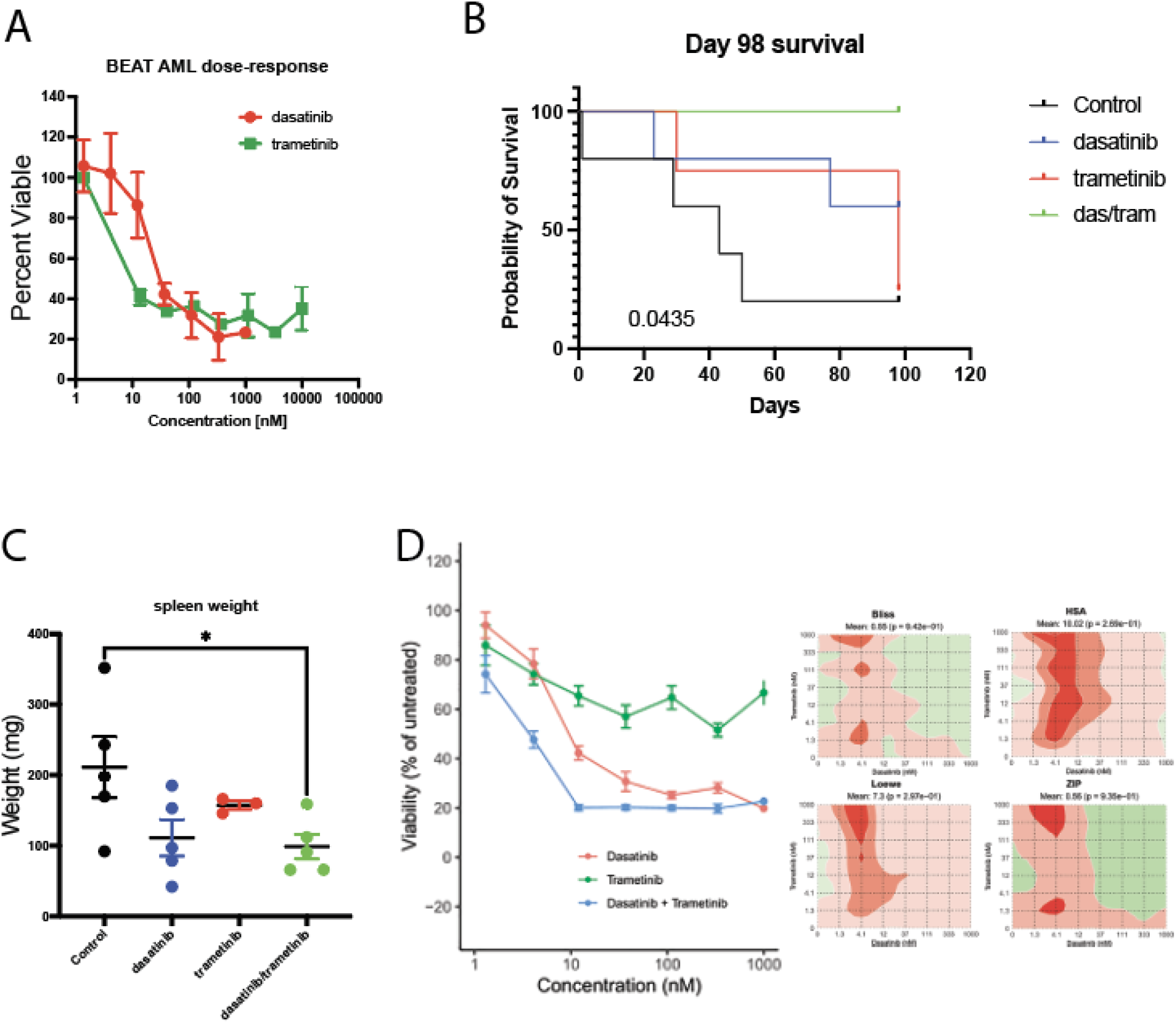
in vivo response to combination therapy in Primary AML PDX B) Primary AML sample with PTPN11 F71L mutation treated with trametinib and dasatinib. A primary AML sample (15-892) carrying a P71L PTPN11 mutation was engrafted into cohorts of five NSGS mice. Upon signs of engraftment with peripheral blood chimerisms of 0.1 to 3% human CD33/CD45, animals were treated with vehicle (Control), dasatinib (40mg/kg/day), trametinib (1mg/kg/day), or combination (40mg/kg/day dasatinib/1mg/kg/day trametinib). Survival wa monitored for either ≥25% human CD33/CD45 in the peripheral blood or the animal becam moribund. Sacrificed animals were assessed for disease status by spleen weight. By day 98 of treatment, remaining animals were sacrificed to evaluate the disease status with the spleen weight. Statistical comparisons were performed with a Log-rank test for survival comparison (p<0.05 comparing control to das/tram) and student’s t-test. * p<0.05.

## Discussion

In these studies, we have validated the utilization of the Ptpn11^D61Y^ mouse model with LysMcre to produce a myeloproliferative neoplasm mirroring JMML. This unique model allowed the focus to lie in the effects of PTPN11 activation in the myeloid lineage. Utilizing this model, we were able to validate combined use of the MEK inhibitor, trametinib, with the TNK2/SRC inhibitor, dasatinib, with *in vivo* therapeutic effect that exceeded responses observed with both single agents. This combination was well tolerated within the mouse for an extended treatment period of three months and the treatment effect with combination included resolution of disease parameters to near wildtype cohorts. Finally, this combination, tested in a patient derived xenograft of AML showed improved treatment effect.

Currently, there is an active interest in therapeutic pursuit of MEK inhibition in JMML (13). Further, single agent MEK inhibition has also shown transient therapeutic effect in RAS mutant AML (32). All prior experience with MEK inhibition has been transient requiring the pursuit of combinations. With strong preclinical evidence the addition of hypomethylating agent in combination is being tested in JMML (NCT05849662). The addition of hypomethylating agents only showed modest activity in AML suggesting alternate approaches will be required (33). The combination of dasatinib and trametinib led to a reduction in disease burden in both a genetic mouse model of JMML as well as a patient derived xenograft of AML carrying a PTPN11 activating mutation. Since both trametinib and dasatinib as single agents and combined with cytotoxic chemotherapy are well tolerated, it is our hope that our findings can rapidly translate to clinical testing.

## Author contributions

BHC conceived, designed, performed the experiments, and co-wrote the manuscript. KTK assisted in the study design and performed experiments. J.W.T. conceived, assisted in the design, and co-wrote the manuscript.

## Data and materials availability

All other data that support the findings of this study are available from the corresponding author upon reasonable request.

